# Analysis of receptor-ligand pairings and distribution of myeloid subpopulations across the animal kingdom reveals neutrophil evolution was facilitated by colony-stimulating factors

**DOI:** 10.1101/2020.06.19.161059

**Authors:** Damilola Pinheiro, Marie-Anne Mahwin, Maria Prendecki, Kevin J Woollard

## Abstract

Neutrophils or heterophils constitute the largest population of phagocytic granulocytes in the blood of mammals and birds. The development and function of neutrophils and monocytes is primarily governed by the granulocyte colony-stimulating factor receptor family (CSF3R/CSF3) and macrophage colony-stimulating factor receptor family (CSF1R/IL34/CSF1) respectively. Using various techniques this study considered how the emergence of receptor:ligand pairings shaped the distribution of blood myeloid cell populations. Comparative gene analysis supported the ancestral pairings of CSF1R/IL34 and CSF3R/CSF3, and the emergence of CSF1 later in tetrapod lineages after the advent of Jawed/Jawless fish. Further analysis suggested that the emergence of CSF3 lead to reorganisation of granulocyte distribution between amphibian and early reptiles. However, the advent of endothermy likely contributed to the dominance of the neutrophil/heterophil in modern-day mammals and birds. In summary, we show that the emergence of CSF3R/CSF3 was a key factor in the subsequent evolution of the modern-day mammalian neutrophil.

**Impact statement:** Colony-stimulating factors (CSFs) are important for myeloid phagocyte development. The emergence of CSF3/CSF3R in tetrapod lineages has uniquely contributed to physical, functional and structural adaptions observed in mammalian neutrophils.

## Introduction

Phagocytes are key effector immune cells responsible for various biological processes; from orchestrating responses against invading pathogens to maintaining tissue homeostasis and neutrophils are the most abundant population of granulocytic phagocytes present in mammalian blood [1–5]. Neutrophil and heterophils (their non-mammalian counterpart) arise from a shared pool of haematopoietic stem cells and mitotic myeloid progenitor cells that can also differentiate into monocytes, eosinophils, and basophils following exposure to the relevant growth factor [1, 4, 6]. The development and life cycle of mammalian neutrophils through a continuum of multipotent progenitors to a post-mitotic mature cell has been well described and has been recently reviewed [2, 4].

The development and function of myeloid phagocytes is mediated through lineage-specific transcription factors and pleiotropic glycoproteins - termed colony-stimulating factors (CSFs)-acting in concert on myeloid progenitor cells. CSF1, CSF3 and their cognate receptors, are lineage-specific and responsible for the differentiation and function of monocytes/macrophages and neutrophils respectively [7–11]. There is a large body of evidence demonstrating the requirement of CSFs for cell development as multiple studies in knockout mice have shown that CSF1R/CSF1 and CSF3R/CSF3 are linked to the development of monocytes and neutrophils *in vivo.* The loss of CSF3 or CSF3R directly affects neutrophil populations resulting in a severe neutropenia, but not the complete loss of mature neutrophils in the models studied [10–12]. The loss of CSF1 caused reduced cavity development of the bone marrow, loss of some progenitor populations, monocytopenia and reduced population of neutrophils in the bone marrow, although interestingly, elevated levels of neutrophils were observed in the periphery [7, 8, 13].

Similarly, in humans, single gene mutations have been described in both CSF3 and CSF3R resulting in severe congenital neutropenia (SCN). In contrast to some animal models, individuals present as children with early onset life-threatening infections because they lack mature neutrophils in the circulation as the neutrophil progenitors in the bone marrow do not progress beyond the myelocyte/promyelocyte stage [14]. These studies demonstrate that CSF receptor/ligand pairings are essential for homeostatic neutrophil development and are intrinsically linked with neutrophil function, arguably making them ideal surrogates to study neutrophil evolution. Through multiple methods we examined the emergence of the respective CSF ligand and receptor genes and proteins across the Chordate phylum and demonstrated how CSF1R/CSF1 and CSF3R/CSF3 pairings contributed to the evolutionary adaptions of the mammalian neutrophil.

## Results

### The neutrophilic/heterophilic granulocyte is the predominant granulocyte in the blood of mammals and aves

The presence of analogous myeloid granulocytes and agranulocytes was examined by comparing available complete blood count (CBC) data, where applicable, for various animal orders and demonstrated the possible distribution of myeloid cells in blood across evolution. The earliest chordates were represented by two groups, the first; jawed fish (4.10 × 10^8^ MYA), which included; coelacanths, elasmobranchs and whale sharks. Jawless fish containing lampreys and hagfish (3.6 × 10^8^ MYA) represented the second group. The next group represented chronologically was the amphibians (3.5 × 10^8^ MYA); containing anurans and gymnophiona. The reptilian class was represented by three different orders, squamata (3.3 × 10^8^ MYA), crocodilia (2.4 × 10^8^ MYA) and testudines (2.1 × 10^8^ MYA). The following group was the closely related avian class (7.0 × 10^7^ MYA), which, contained birds from both paleognaths and neognaths. The final class of chordates evaluated was Mammalia, where all three orders; monotremata (1.1 × 10^7^ MYA), marsupial (6.5 × 10^7^ MYA) and placental (6.25 × 10^7^ MYA) were represented. All the species examined for cell distribution, and for the subsequent gene sequence and/or protein homology studies were visualised in a species tree (Figure 1a).

**Figure 1.**
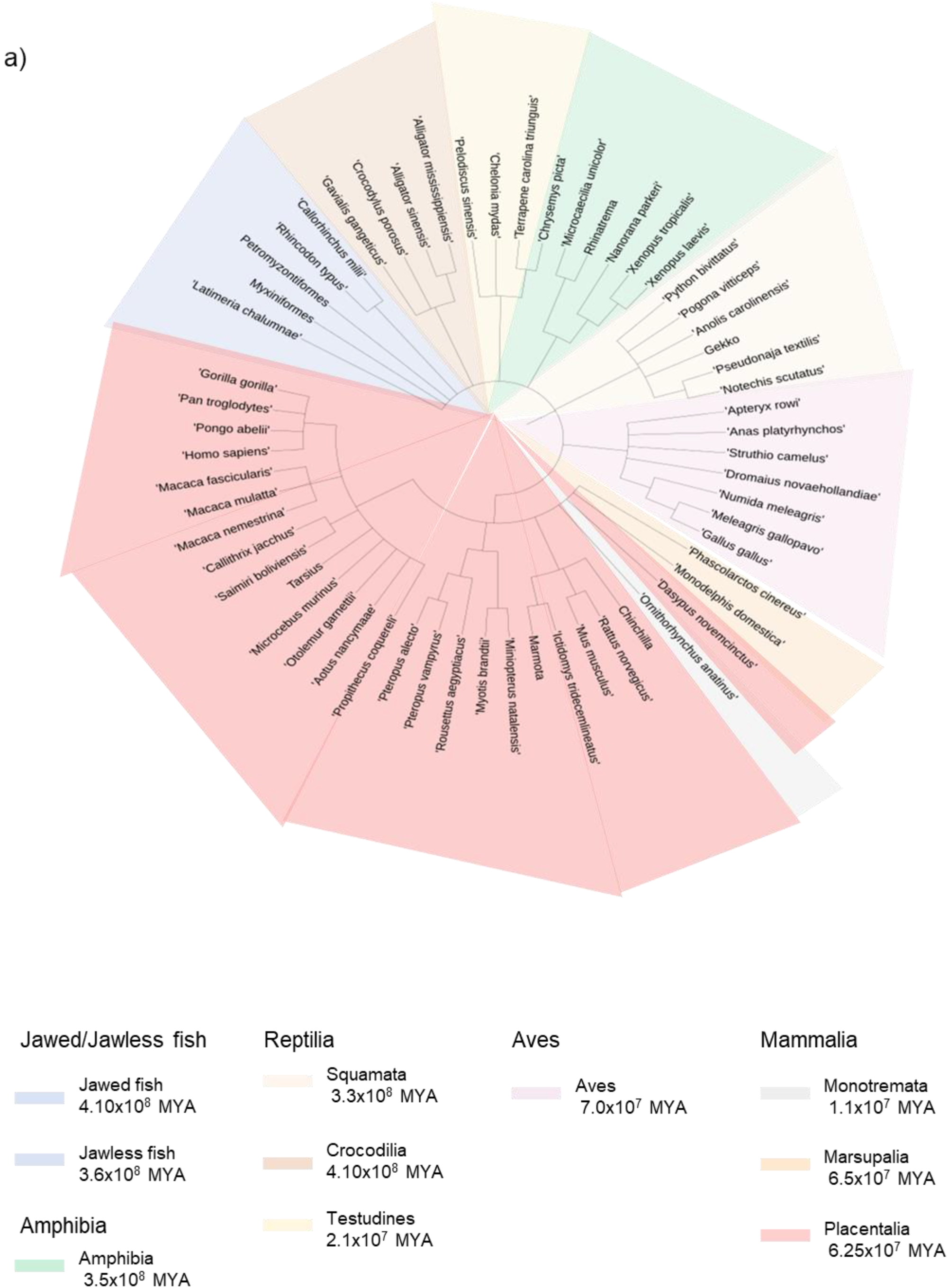
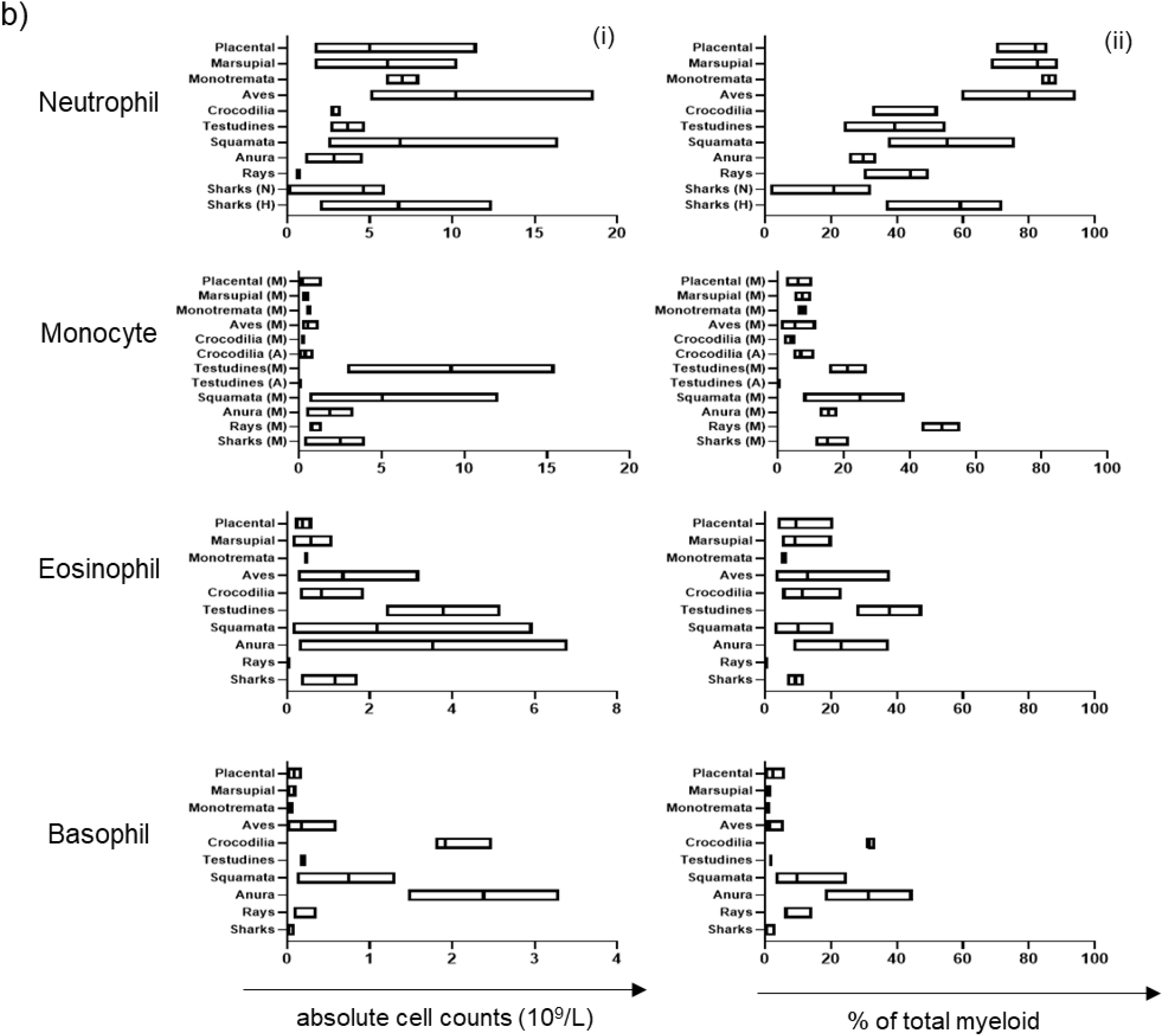
Population comparison of blood myeloid cell subset distribution in chordates demonstrates predominance of the neutrophilic granulocyte in birds and mammals. A. Species tree of the animals within Phylum Chordata examined, sub-classified by animal order or class. B. Meta-analysis of aggregated phylum Chordata complete blood cell counts (excluding all lymphoid cells). Data is visualised as a floating bar and the line represents the mean value and shows absolute counts per cell type per order (i) and composition of myeloid cells per cell type (ii).

All classes within the phylum Chordata were evaluated for populations of neutrophils/heterophils; eosinophils and basophils. Azurophils, a specialised population of granulocytes – analogous to both the mammalian neutrophil and monocyte-but unique to reptiles were also included. A meta-analysis was performed on aggregated data to show absolute counts per cell type per order and finally, the proportional composition of myeloid cells only per cell type (Figure 1b).

The presence of all myeloid cells was confirmed in at least one representative species from each order examined. Elasmobranchs-rays and sharks-were unique in having both heterophils (2.009 – 12.42 ×10^9^, n=3) and neutrophils (0.615 – 0.745 ×10^9^, n=3). All other orders had either neutrophilic or heterophilic populations only. Heterophils were the predominant granulocyte in Amphibians (the order of Anura; 1.10 – 4.60 ×10^9^, n=2), Reptiles (the orders; squamata, 2.51 – 16.43 ×10^9^, n=6; testudines, 2.59 – 4.74 ×10^9^, n=2; and crocodiles, 2.59 – 3.24 ×10^9^, n=3) and Aves (0.615 – 0.745 ×10^9^, n=5). In contrast, neutrophils were the majority population in all three mammalian orders (monotremata, 5.97 – 8.01 ×10^9^, n=2; marsupalia, 1.64 – 10.30 ×10^9^, n=4; and placental, 2.04 – 11.50 ×10^9^, n=5 [Figure 1bi]). Eosinophils and basophils were over-represented in the orders of amphibians, reptiles and aves, when compared to mammals, where they were either a minority population (eosinophils) or absent (basophils) [Figure 1bi].

Analysis of the proportional composition of total myeloid white blood cells (WBC) showed interesting changes in the distribution of granulocytes across the different species classes. Again, elasmobranchs were unique amongst all classes as they retained both a majority heterophil population (36.8% – 71.9%, n=3) and a minority neutrophil population (1.6% – 32.3%, n=3). The cold-blooded orders; rays (30.0% – 39.8%, n=3), anura (25.6% – 33.7%, n=2), squamata (37.3% – 75.7%, n=6), testudines (23.9% – 54.7%, n=2) and crocodilia (32.4% – 52.4%, n=3); retained a large population of neutrophils/heterophils; although proportions of the population varied within related animal orders and between the different species classes (Figure 1bii). In contrast, within the warm-blooded classes and orders; aves (59.6% – 94.3%, n=5), monotremata (83.8% – 88.4%, n=2), marsupial (68.5% – 88.8%, n=4) and placental (70.0% – 85.6%, n=5), the opposite distribution was observed and the neutrophil/heterophil was the most abundant peripheral blood cell type (Figure 1bii). Interestingly, basophils and eosinophils were well represented in the anuran order, 18.3% – 44.8%, n=2, and 8.6% – 37.7%, n=2, respectively. A similar pattern was observed in the reptilian lineage; however, there were differences between orders as squamata (2.9% – 20.8%, n=6) and crocodilia (5.3% – 23.4%, n=3) had lower proportions of eosinophils compared to testudines (27.7% – 47.6%, n=2). Conversely, basophils were elevated in crocodilia (31.1% – 33.6%, n=3) compared to squamata and testudines (3.4% – 24.9%, n=6 and 1.2% – 2.19%, n=2 respectively) [Figure 1bii]. Suggesting that within closely related orders that share similar habitats or environments, individual species favoured different configurations of myeloid WBCs. Eosinophils were present as a small population in the mammalian lineage, again variation between the respective classes was observed as monotremata had a smaller proportion of cells (4.9% – 6.7%, n=2) compared to the marsupial and placental orders, which had similar values (5.1% – 20.3%, n=4 and 3.8% – 20.8%, n=5 respectively). In contrast, compared to all the other classes, basophils seemed to have been largely lost from some of the mammalian classes, as low proportions were observed in monotremata (0.6% – 0.6%, n=2), marsupial ((0% – 1.85%, n=4) and placental classes (0% – 6.2%, n=5) [Figure 1bii].

Monocytes are blood agranulocytic cells that are closely related to neutrophils sharing both growth and survival factors, early development pathways with common progenitors, and in some instances, effector functions in the respective terminally differentiated cells [5, 15, 16]. Interestingly, the monocyte population peaked both in terms of number and proportion within the reptilian class, in the orders; squamata (0.6 – 12.1 ×10^9^, n=6 and 15.4% – 38.5%, n=6 respectively) and testudines (2.9 – 15.5 ×10^9^, n=2 and 7.8% – 27.0%, n=2 respectively). The azurophil – a specialised myeloid cell population-peaked in the crocodilian lineage (4.6 – 11.1 ×10^9^, n=3 and 0.2% – 0.9%, n=3 respectively), however; it remains unclear whether to classify the azurophil as a distinct cell type, as we have done, or as a subset of monocyte or neutrophil. Aves is the most closely related phylogenetic group to reptiles as they share a recent common ancestor. Although Aves has lost the azurophil, they retain a small population of monocytes (2.4% – 11.7%, n=5). Interestingly, a similar proportion is observed in the more distantly-related mammalian class, across all three orders (monotremata (6.2% – 8.8%, n=2), marsupial (5.1% – 10.2%, n=4) and placental (2.5% – 10.4%, n=5), suggesting that the advent of endothermy may have played a role in the distribution of monocytes in warm-blooded animals [Figure 1bii] In summary, we show that the majority of peripheral myeloid blood is comprised of granulocytes in phylum Chordata, although the proportion of basophils, eosinophils and neutrophils varies according to the respective orders and lineages. By the advent of mammals and birds however, the neutrophil has become the predominant granulocyte of the blood.

### Interrogation of CSF1/CSF1R and CSF3/CSF3R and C/EBP gene family reveals their loss in the early lineages

The development and maturation of a neutrophil from a multipotent progenitor to a lineage committed post-mitotic cell is driven by CSF3 working in concert with a number of transcription factors, including Pu.1, GF1I and Runx, whose roles have been well established [4, 16]. One of the most heavily involved family of transcription factors in neutrophil development is the CCAT enhancer binding-protein family (C/EBP), comprising six members (C/EBPα, β, δ, γ, ε and τ) [16–19]. Individual members are a necessary requirement for the different stages of granulopoiesis under homeostatic and inflammatory conditions. Studies have shown that *c/ebpα*^*−/−*^ mice are neutropenic, whereas *c/ebpε*^*−/−*^ mice lack functionally mature neutrophils, thus demonstrating that *c/ebpα* and *c/ebpε* have distinct roles in the early (myeloid progenitor progression) and later (maturation) stages of neutrophil development [16–19]. Having established, the presence of the neutrophil/heterophils across the phylum Chordata, we then evaluated the presence of CSFR/CSF and C/EBP genes in the different animal classes to further understand the emergence of granulopoiesis in phylum Chordata.

Gene data for fifty-nine species was collected from the NCBI Gene and Ensembl databases for the following genes; CSF1R, IL34, CSF1, CSF3R, CSF3, C/EBPα, C/EBPβ C/EBPδ and C/EBPα and used to generate a heat map. Interestingly, CSF1R, CSF3R and CSF3 are absent from the cartilaginous fishes; the Australian Ghost shark and the Whale shark (Figure 2a). In contrast, the Coelacanth - another early lineage -retained CSF1R, IL34, CSF3R and CSF3 genes. CSF1 was absent from all the jawed fishes although IL34 was present. A similar pattern was observed in the Jawless fish, CSF1R, IL34 and CSF3R were present in the Hagfish, whereas CSF1 and CSF3 were absent. CSF1R was the only the receptor present in the lamprey and all other CSF receptor and ligands were absent (Figure 2a). The Jawed/Jawless fish accounted for the largest loss of genes within in an animal order as generally; CSF1R, IL34, CSF1, CSF3R and CSF3 were largely all present in amphibia, reptilia, aves and mammals. The exception being in the reptilian order of squamata, where IL34 had been lost. There were also examples of gene loss at the species level, CSF1 was absent from the Gharial and Ostrich; however, it had not been lost overall in the respective orders of crocodilia and struthioniformes (Figure 2a).

**Figure 2.**
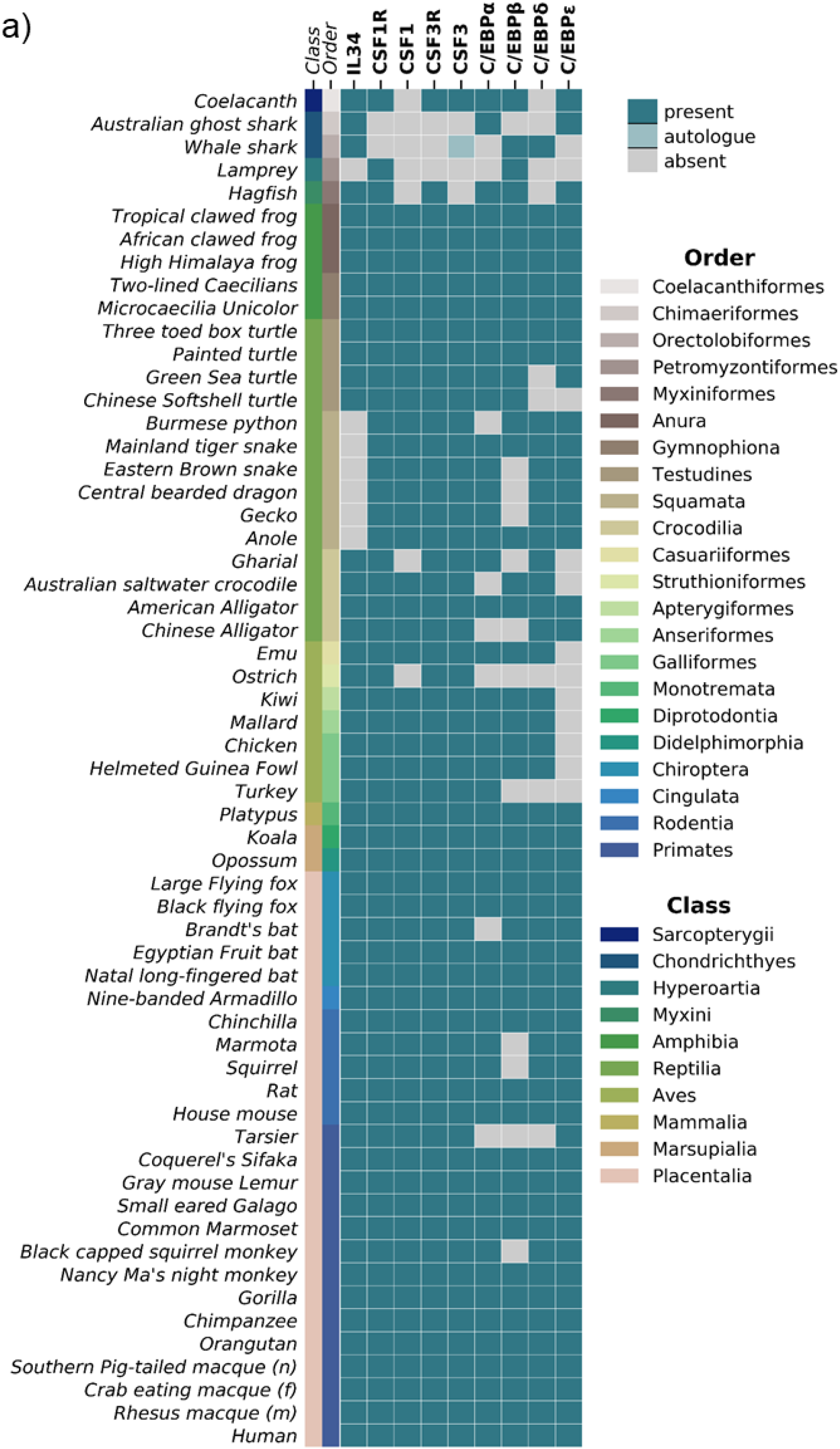
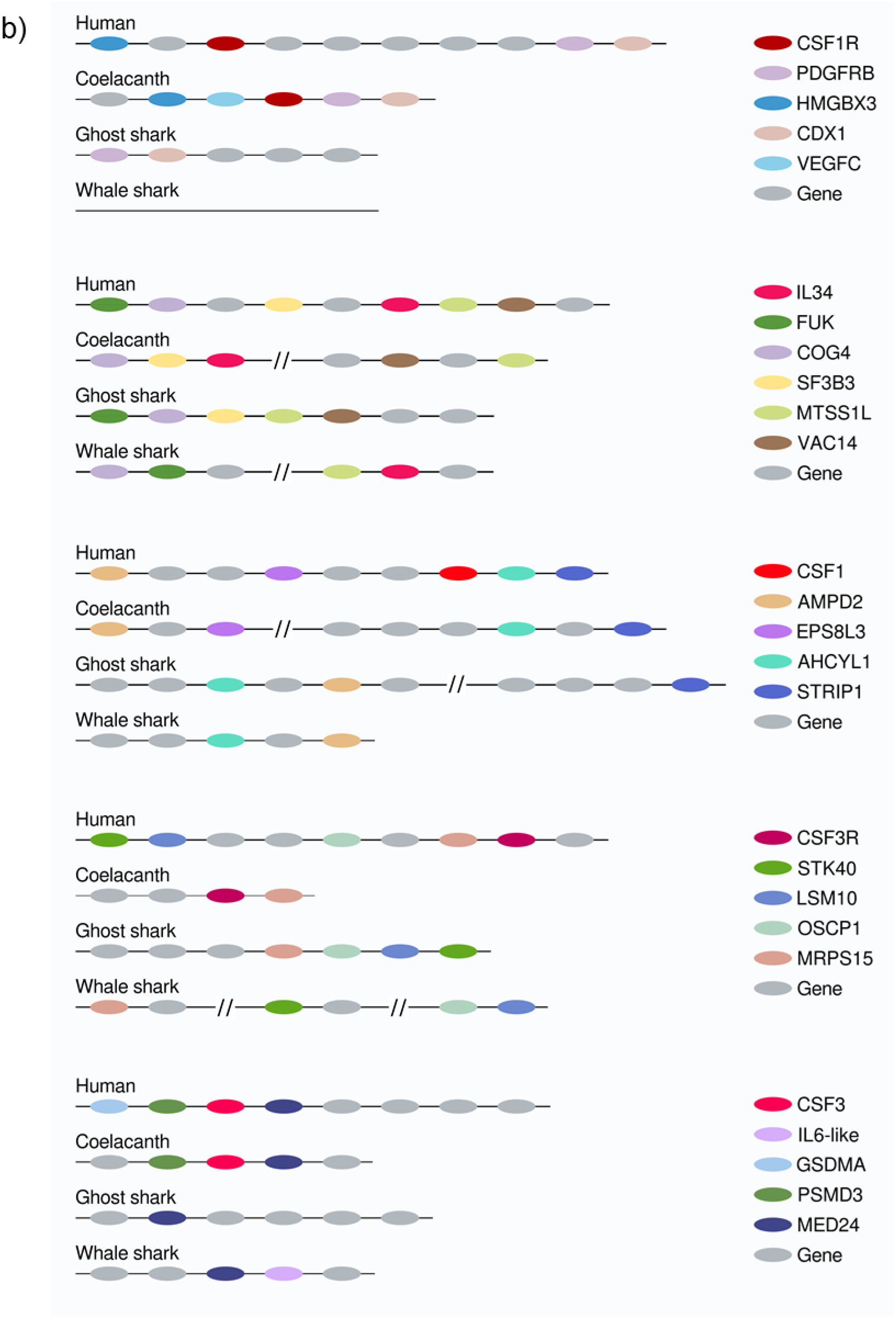
Analysis of chordate CSF1/CSF1R, CSF3/CSF3R and neutrophil-related transcription factors reveals their loss in Jawed/Jawless fish lineages. A. Heat map of CSF1R/IL34/CSF1, CSF3R/CSF3 and neutrophil-related transcription factors in selected members of Phylum Chordata. B. Syntenic maps of CSF1R/IL34/CSF1 and CSF3R/CSF3 in selected Jawed fish compared to human.

The Jawed/Jawless fishes were again the most diverse in terms of coverage within the C/EBP gene family. The Coelacanth had present all members examined (C/EBPα, C/EBPβ, C/EBPδ and C/EBPε), the Hagfish had three members (C/EBPα, C/EBPβ, and C/EBPε) and the Australian Ghost shark (C/EBPα and C/EBPε) and the Whale shark (C/EBPβ and C/EBPδ) had two members each. Finally, the Lamprey had only one C/EBP gene present, C/EBPβ (Figure 2a). Within the other animal orders, C/EBP gene loss was largely restricted to individual species, such as the loss of C/EBPβ in the Eastern Brown snake, Central Bearded dragon and Gecko; however, it was not lost across the whole order of Squamata. In the order of testudines, both C/EBPδ and C/EBPε or C/EBPε only, had been lost from the Green Sea Turtle and Chinese Softshell turtle respectively (Figure 2a). Similarly, in the Crocodilian order, the Gharial (C/EBPδ and C/EBPε), Australian Saltwater Crocodile (C/EBPδ and C/EBPε), and Chinese Alligator (C/EBPδ and C/EBPε) all had two different C/EBP genes present, in contrast to the American alligator, which retained the complete set (C/EBPα, C/EBPβ, C/EBPδ and C/EBPε). Interestingly, while the C/EBPε gene was missing from all birds examined, the ostrich appeared to have no C/EBP gene family members present and the turkey only retained C/EBPα (Figure 2a). Finally, there was some individual species-specific loss of C/EBP genes in the mammalian class, with examples of C/EBPα and C/EBPβ loss being observed in some members (Figure 2a). C/EBPε, which is required for the later stages of mammalian neutrophil development was present across the order (Figure 2a). These results suggested that there were local factors determining the retention and loss of genes as well as a strong level of functional redundancy among the C/EBP gene family.

Interestingly, within the early lineages of the jawed fishes, IL34 was the only gene retained by all representatives and the Coelacanth was the only species to have CSF1R, CSF3R and CSF3 present. Given that these species all share a close common ancestor, this would indicate that there had been local gene loss in both the Whale shark and Australian Ghost shark. CSF1 was absent from all lineages and it was unclear as to whether this absence was due to loss or that CSF1 had not yet evolved. To further address this, we used syntenic methods to manually map out the orthologous gene locations in the Jawed fish. Syntenic maps were generated for the CSF1R (CSF1R, IL34, CSF1) and CSF3R (CSF3R and CSF3) family of genes using the human orthologue as a reference point (Figure 2b).

In humans, the CSF1R gene is located downstream of the HMGBX3 gene and upstream of PGDFRB (a gene paralogue of CSF1R) and CDX1. A similar orientation is observed in the Coelacanth, where CSF1R is downstream of HMGBX3 and adjacent to PDGFRB and CDX1. The orientation is inversed in the Australian Ghost shark, where PDGFRB and CDX1 colocalise together upstream but HMGBX3 and CSF1R have been lost (Figure 2b). Interestingly in the Whale shark, CSF1R and all the flanking genes are absent, suggesting that this section in its entirety may have been lost (Figure 2b). Human IL34 co-localises with FUK, COG4 and SF3B3 upstream, and the MTSS1L and VAC14 genes immediately downstream. Both the Coelacanth and the Whale shark have partially retained the syntenic combination but not in the same location. IL34 is situated immediately downstream of COG4 and SF3B3, whereas MTSS1L and VAC14 are located elsewhere on the Coelacanth chromosome (Figure 2b). In the Whale shark, IL34 is immediately adjacent to MTSS1L and these genes are both downstream of FUK and COG4, which have co-localised next to each other (Figure 2b). The gene arrangement of the Australian ghost shark most closely resembles the human, as FUK, COG4, SF3B3, MTSS1L and VAC14 all co-localise to the same region on the chromosome and the absence of IL34, suggest it has been loss in the process of a local gene rearrangement (Figure 2b). The CSF1 human orthologue has formed a contiguous block with AHCYL1 and STRIP1 and is flanked upstream by AMPD2 and EPS8L3. Interestingly although CSF1 is absent from all members, the flanking genes have largely been retained. Similar to the human arrangement, in the Coelacanth; AMPD2 and EPS8L3 co-localised together, while AHCYL1 and STRIP1 are located in close proximity in a different location (Figure 2b). In both the Australian Ghost and Whale Sharks, AHCYL1 and AMPD2 are located together, STRIP1 is located elsewhere in the Australian Ghost shark and has been lost entirely from the Whale shark (Figure 2b). These lineages arose early in evolution but have very similar synteny structure to the later emerging human chromosome, suggesting that the CSF1 gene entered this location in an ancestor that emerged after the jawed fish.

A similar pattern emerged in the CSF3 family of proteins when analysed. In humans, the CSF3R gene is located downstream of STK40, LSM10, OSCP1 and MRPS15. The Coelacanth is the only species to retain CSF3R and that is located immediately adjacent to MRPS15, however STK40, LSM10 and OSCP1 have been lost. In contrast, although both the Whale shark and Australian Ghost shark have presumably lost the CSF3R gene, they have retained the four other flanking genes, either in one location as in the Australian Ghost shark, or distributed along the chromosome, as in the Whale Shark (Figure 2b). Human CSF3 co-localises downstream of GSDMA and PSMD3 and is adjacent to MED24. Again, a similar arrangement is observed in the Coelacanth, with CSF3 flanked by PSMD3 and MED24 upstream and downstream respectively. As before, the Whale shark and Australian Ghost shark are similar in their gene arrangements as GSDMA and PSMD3 have been lost, and the only gene retained in the Australian Ghost shark is MED24 (Figure 2b). Intriguingly, in the Whale shark, MED24 co-localises with an IL6-like gene, which could be a functional paralogue of CSF3 (Figure 2b). These results suggest there is more than one receptor ligand family involved in the development and maturation of heterophils and neutrophils in the early lineages. Taken together, these data suggest that CSF3R/CSF3 and CSF1R were present in a common ancestor to early lineages and there have been so local losses in selected members. CSF1 appears to have evolved independently of its cognate receptor, after the emergence of Jawed/ lineages and prior to the advent of the tetrapod lineages.

### Analysis of Chordate orthologous protein homology further supports the ancestral pairing of CSF1R/IL34 and CSF3R/CSF3 in early lineages

The syntenic analysis established the presence of the CSF1R and CSF3R gene families in Chordates. Interestingly, CSF1R/IL34 and CSF3R/CSF3 had already evolved by the emergence of Chordates, as evidenced by their existence in some of the early lineages of Jawed/Jawless fish. However, the gene data in isolation did not provide a complete understanding and additional analysis was needed. Orthologous proteins are considered to have the same function in different species and therefore, it’s broadly assumed that the proteins will be largely conserved at the primary and structural levels [20]. To further elucidate the evolutionary process, we compared the shared sequence similarity of the orthologous CSF1R and CSF3R protein families, as this data is widely available. The protein sequences for the following proteins; CSF1R, IL34, CSF1, CSF3R and CSF3 for fifty-nine species were collated from the NCBI Protein database. The shared sequence similarity for each individual sequence was generated using the NCBI BLAST engine tool and the relevant human orthologue submitted as the query. The resulting data was then plotted in groups as identified by animal order and visualised in a bar chart (Figure 3a).

**Figure 3.**
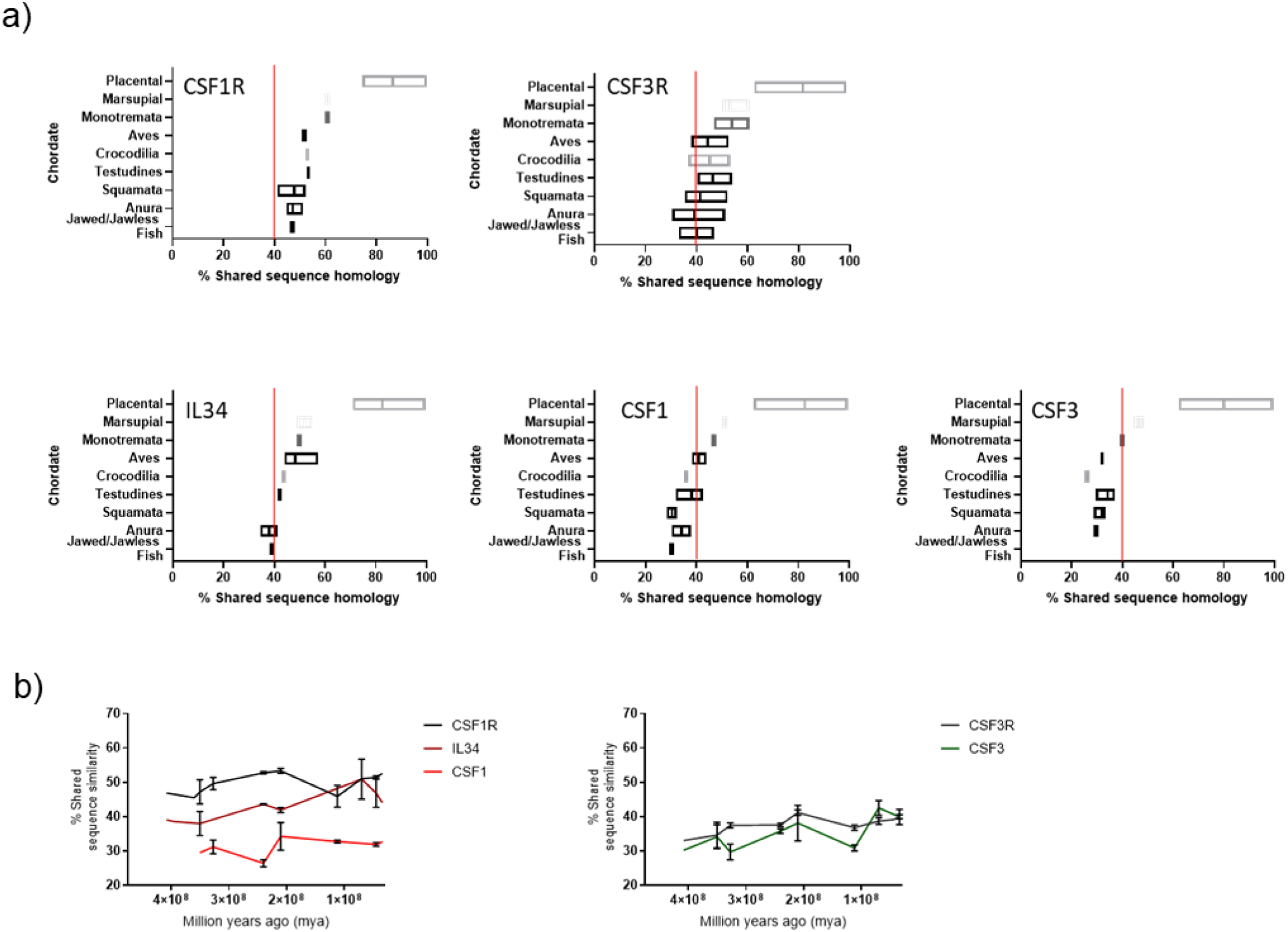
Shared sequence similarity analysis of Chordate CSFR/CSF protein homology further supports the ancestral pairing of CSF1R/IL34 and CSF3R/CSF3 in early lineages. A. Representative plots of % shared sequence similarity for CSF1R, CSF3R, IL34, CSF1 and CSF3 in the respective sub-groups of Phylum Chordata, data is visualised as a floating bar and the line represents the mean value. B. Graphical representation of % shared sequence similarity in sub-groups of Phylum Chordata *versus* time.

The level of shared sequence similarity observed in the receptors, and therefore conservation of the primary protein sequence, varied across the different animal orders. The highest levels of shared sequence conservation for both CSF1R and CSF3R were observed in the placental mammals (CSF1R; 86.5%, CSF3R; 81.6), which would be anticipated as the human is a member of this group. The lowest level of shared sequence similarity was observed in the Jawed/Jawless fish (CSF1R; 46.9%, CSF3R; 40.0%) (Figure 3a). Interestingly, the CSF1R and CSF3R protein sequence values from early mammals, monotremata (CSF1R; 60.6%, CSF3R; 53.8%) and marsupalia (CSF1R; 60.8%, CSF3R; 53.1%), ranged between the earlier orders; Aves (CSF1R; 51.4%, CSF3R; 44.4%), Testudines (CSF1R; 53.3%, CSF3R; 46.4%), Crocodilia (CSF1R; 52.8%, CSF3R; 45.2%), Squamata (CSF1R; 47.8%, CSF3R; 41.5%), Amphibia (CSF1R; 47.2%, CSF3R; 39.4%) and placental mammals (Figure 3a). While there is not a clear consensus as to what the minimum percentage of shared sequence similarity needed to correlate with conservation of function is, it is notable that in this dataset for both CSF1R and CSF3R - independently of each other-the baseline value of shared sequence similarity was approximately 40% (Figure 3a).

The syntenic analysis also demonstrated that of the three ligands, IL34 emerged the earliest and this was reflected in the protein data. As observed with the receptors, the highest level of shared sequence similarity is in the placental mammal group (82.5%), and the lowest in the Jawed/Jawless fish (38.8%) (Figure 3a). The IL34 protein sequence values from early mammals; monotremata (50.0%) and marsupalia (50.2%), grouped very closely with the earlier orders of; Aves (48.2%), Crocodilia (43.6%), Testudines (41.9%), Amphibia (38.0%) and the Jawed/Jawless fishes (Figure 3a). IL34 was absent from the members of Squamata examined. CSF3 is the next best conserved ligand, which again agreed with the synteny data. The highest level of shared sequence similarity was in the placental mammal group (82.6%), and the lowest in the Jawed/Jawless fish (30.2%) (Figure 3a). The CSF3 protein sequence values were spread among the orders from monotremata (46.9%) and marsupalia (51.2%) to Aves (41.0%), Crocodilia (35.8%), Testudines (38.2%), Squamata (30.5%) and Amphibia (34.1%) (Figure 3a). Finally, CSF1, which is completely absent in Jawed/Jawless fish, had similar values to CSF3, in terms of shared sequence similarity across the respective groups of; placental mammals (80.0%), monotremata (46.5%), marsupalia (40.0), aves (32.2%), Crocodilia (26.5%), Testudines (34.3%), Squamata (32.1%) and Amphibia (29.5%). Intriguingly, the 40% baseline is applicable to the ligand data. In IL34, which is the oldest ligand, the shared sequence similarity for all groups was above 40%. Whereas for both CSF3 and CSF1, which emerged later, in the majority of early groups the shared sequence similarity was under 40% (Figure 3a).

The analysis of orthologous CSF1R and CSF3R protein families illustrated that the mammalian proteins-largely within the placental mammals-had changed considerably compared to the other lineages and therefore were excluded from subsequent analysis. To further interrogate how the respective CSF1R and CSF3R families co-evolved in the early lineages, the shared sequence similarities for each order were plotted against time (Figure 3b). Interestingly, the trajectories for CSF1R and IL34 and CSF3R and CSF3, largely tracked to each other. This suggested that they evolved at a similar pace across the same period of time and is indicative of evolutionary pressure restricting changes to cognate receptors and ligands. As expected, CSF1 did not have the same restrictive pattern in the earlier lineages as it emerged much later (Figure 3b). These results supported the early emergence of CSF3R, CSF1R and IL34 of the respective ligands and receptors. Interestingly, although CSF3 developed later than CSF3R, the two have co-evolved in step together. In contrast, while CSF1 would appear to be the principal ligand of CSF1R in mammals, the data supported the ancestral pairing of IL34 and CSF1R in early lineages.

### The emergence of CSF3R/CSF3 and onset of endothermy likely influenced the distribution of neutrophils in Chordates during evolution

The mammalian neutrophil shares many developmental and functional properties with counterparts present in other animal orders such as phagocytosis, oxidative burst, degranulation and cell motility [21, 22]. However; a notable species-specific difference is the of distribution of the neutrophil/heterophil within peripheral blood, which suggests that evolutionary pressures might also be involved. To further address this, we reconstructed a set of related timescales plotting the percentage distribution of chordate myeloid WBC and the percentage of protein shared sequence similarity of chordate CSFR/CSF *versus* time. To simplify the model, the timescales were plotted chronologically, where applicable, and the emergence point of the earliest member within an animal order was used. The timescales were then divided into two distinct periods; the first focussed on the early lineages of Jawed/ Jawless fish, Amphibia and early reptilia and the second focussed on all reptilia, aves and mammalia. We then considered the possible impact the emergence of CSF3 had on cell distribution across the evolutionary timescale (Figure 4).

**Figure 4.**
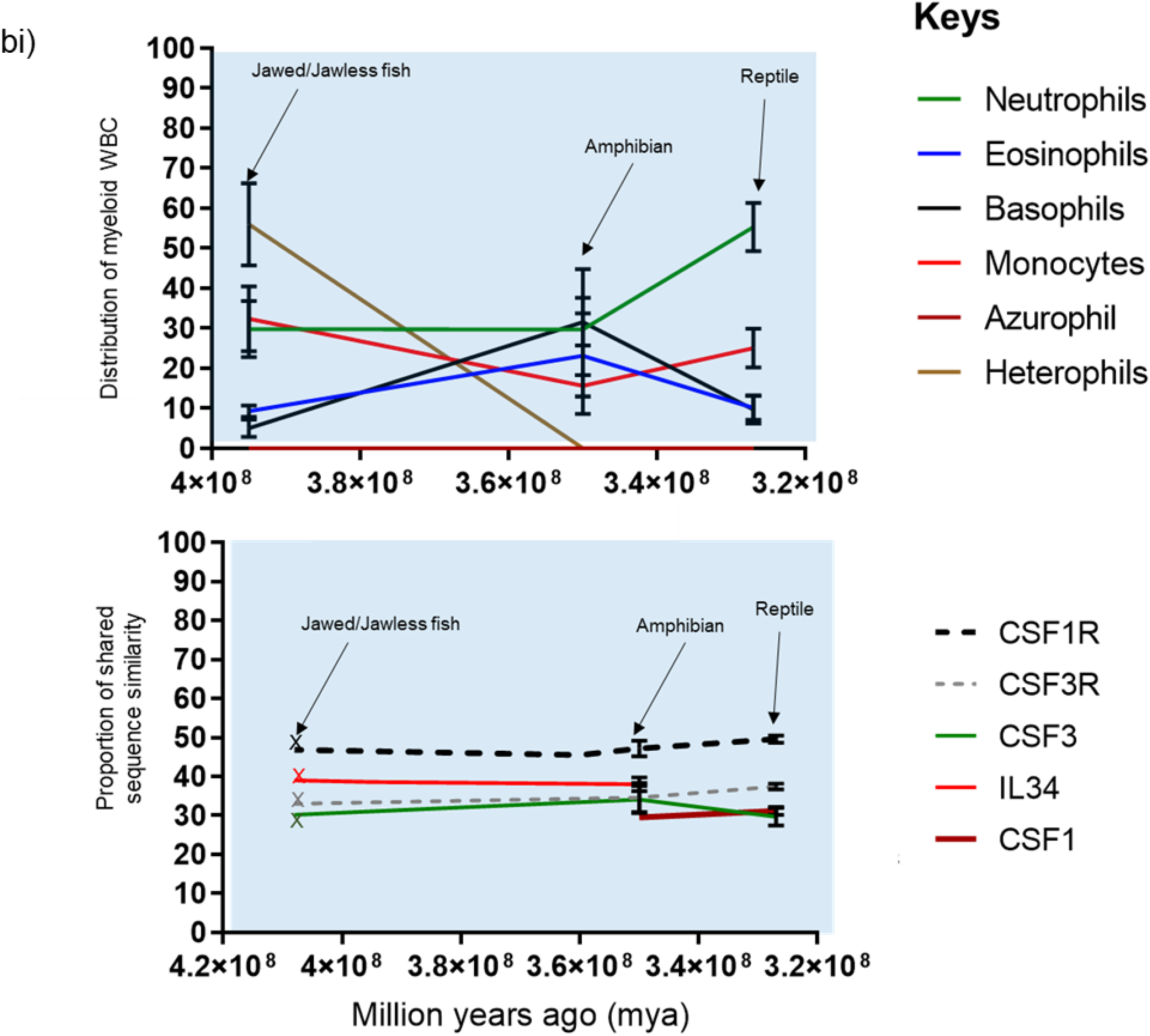
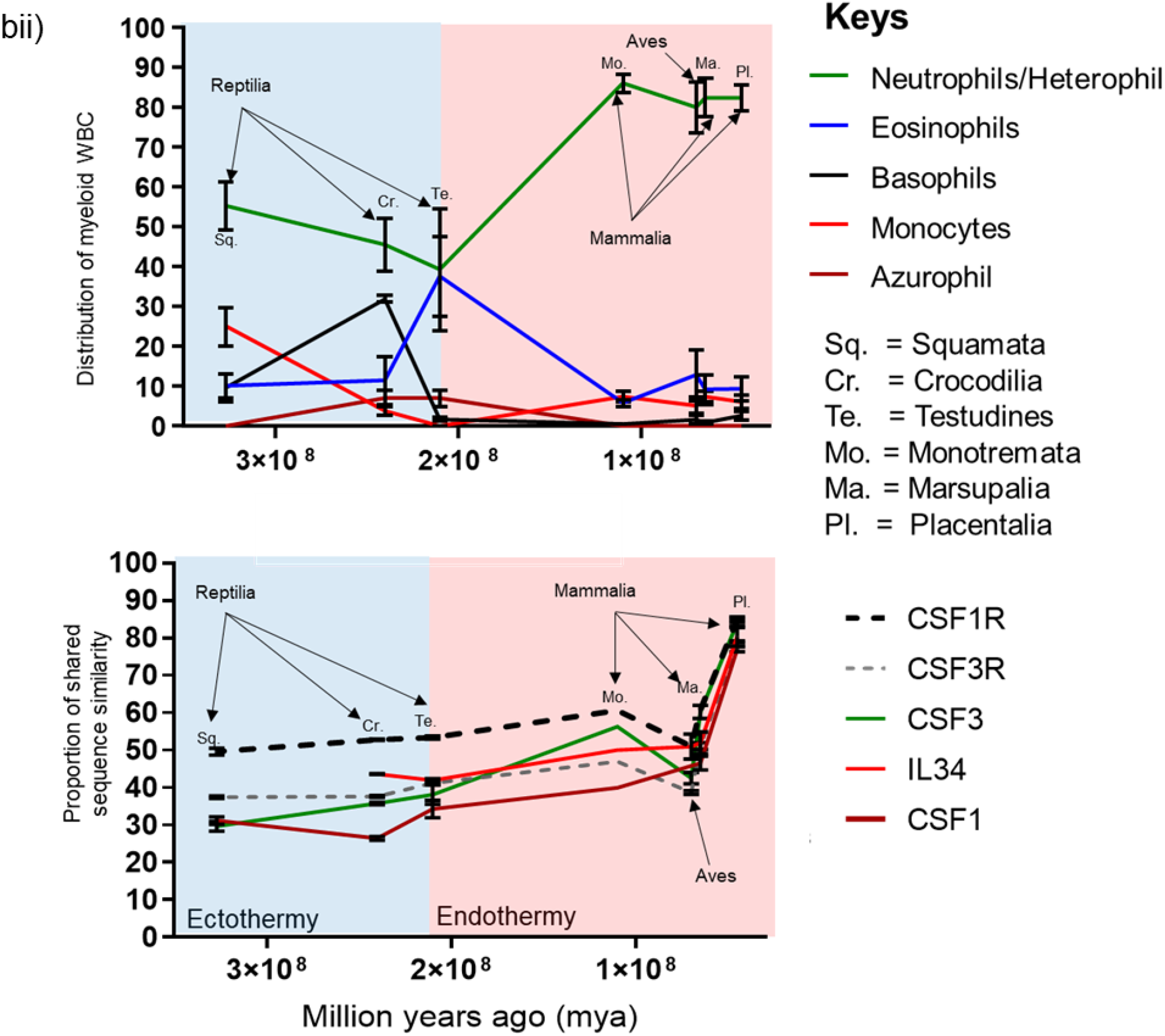
Changes in Chordate blood granulocyte distribution from Jawed/Jawless lineages through to placental mammals are multi-factorial and likely driven by the emergence of CSF3 and onset of endothermy. A. Graphical representation of % population distribution of myeloid white blood cells *versus* time (i) and % shared sequence similarity of CSF1R and CSF3R protein families *versus* time for Jawed/Jawless fish, Amphibia and the reptilian order of Squamata (ii). B. Graphical representation of % population distribution of myeloid white blood cells *versus* time (i) and % shared sequence similarity of CSF1R and CSF3R protein families *versus* time for the Reptilian orders of squamata, testudines and crocodilia, Aves, and the Mammalian orders of monotremata, marsupalia and Placental (ii).

The synteny studies demonstrated that CSF1R and IL34 were already in existence prior to the appearance of the Jawed/Jawless fish lineages. However, CSF3R and CSF3 were only present in the Coelacanth and although an IL6-like paralogue was observed in the Whale shark, a cognate receptor was not identified. These results suggest a greater level of diversity in the growth factors responsible for WBCs in these lineages and is reflected in the distribution of WBCs in Jawed/jawless fish, where heterophils, neutrophils, monocytes, eosinophils and basophils are all present. While heterophils largely dominate, there was a more even distribution of neutrophils and monocytes, with minority populations of basophils and eosinophils (Figure 4ai). By the advent of amphibia, distinct neutrophil and heterophil populations had seemingly been lost in favour of a single neutrophil or heterophil population. The amphibian order was the only lineage to demonstrate a largely even distribution of neutrophils, eosinophils, basophils and monocytes, although there was a slight decline compared to the levels observed in Jawed/Jawless fish (Figure 4ab).

At some stage during tetrapod evolution both CSF3 and CSF1 emerged and haematopoiesis largely transitioned to the tissue-specific compartment of bone-marrow, which likely had implications for WBC distribution in subsequent orders. In early reptilia, represented by the squamata order of lizards, the neutrophil becomes the dominant granulocyte and there is concomitant reduction in basophils and eosinophils. Interestingly, there is an increase in monocyte populations, which coincides with the emergence of the monocyte/macrophage specific growth factor CSF1 (Figure 4ai). These changes occur in ectothermic species suggesting that this is more in response to cell-intrinsic factors, rather than external factors such as environment (Figure 4ai).

We hypothesised that the emergence of CSF1, CSF3 and CSF3R could be one of the cell intrinsic factors as it facilitated the transition of haematopoiesis to the bone marrow through modulating cell mobility. CSF3 and CSF3R in particular, are known to play prominent roles in co-ordinating cell movement out of the bone marrow by disrupting the respective [C-X-C chemokine receptor type 4 (CXCR4) / [C-X-C] motif ligand 12 (CXCL12) retention and CXCR2/CXCL2 egress axes [1, 23, 24]. Although, CSF1R is mostly expressed by monocytes and CSF3R by neutrophils [7, 10], multiple studies have shown that there is receptor cross expression between them. Classical monocytes, the predominant monocyte population in humans, express low levels of CSF3R protein and can be mobilised from the bone morrow in response to exogenous CSF3 [25–27]. Similarly, CSF1R transgene expression has been previously described both in *ex vivo* and cultured murine neutrophils [28]. We hypothesized that the cross expression could be part of an evolutionary conserved mechanism that allowed myeloid cells to modulate the speed of their movement and examined this further in preliminary *in vitro* studies (supplementary Figure 1). Although there was low baseline CSF3R expression (supplementary Figure 1a, monocytes that were cultured in CSF3 for 72 hours moved at speeds comparable to that of freshly isolated neutrophils in *in vitro* mobility assays (supplementary Figure 1b). Conversely, monocytes that were cultured in CSF1 moved the slowest of the populations examined, suggesting that CSF1 and CSF3 signalling pathways have contradictory effects on myeloid cell velocity (supplementary Figure 1b).

Intriguingly, we identified CSF3R expression in a minority population of CXCR4^hi^ neutrophils and negligible expression was observed on the CXCR4^int^ and CXCR4^lo^ populations ex *vivo* (supplementary Figure 1c). CXCR4 expression is upregulated on neutrophils when they home to the bone marrow as well as on populations that are retained in the bone marrow [23, 24]-a microenvironment in which presumably minimal cell movement is required. Therefore, we hypothesised that if this was an intrinsic property of the cell, CSF1R expression would be upregulated and CSF3R downregulated on CXCR4^hi^ neutrophils generated *in vitro*. Neutrophils were cultured in media supplemented with 10% Foetal bovine serum (FBS) for 24 hours, which induced expression of CXCR4 and exogenous CSF3 (20ng/ml) was added to a separate control stream (supplementary Figure 1d). In all cultures, distinct CXCR4^lo^ and CXCR4^hi^ populations were present after 24 hours and CSF1R was only observed in the CXCR4^hi^ populations with a concomitant reduction in CXCR2 and CSF3R (supplementary Figure 1d). The reason for the induction of CSF1R by this subset of neutrophils remains unclear, however, it appears to be cell intrinsic as the presence of exogenous CSF3 had no effect on the induction of expression. While more experimental data is needed, our preliminary in-vitro studies taken together suggest that the ability of neutrophils and monocytes to adjust their velocity, potentially through switching of CSFR expression may have conferred a functional adaption that then facilitated the increase of monocyte and neutrophil populations during the transition from amphibia to early reptiles.

The reptilian lineage comprised three orders; squamata, crocodilia and testudines, who emerged over a large timescale. Accordingly, there are many intra-order differences such as the environments in which members live; i.e. in the water *versus* on land or the presence or absence of limbs or exoskeletons. Interestingly, the neutrophils seemingly peaked in Squamata as proportions steadily declined with the lowest levels observed in Testudines (Figure 4bi). In contrast, Eosinophil levels, which remained low across the squamata and crocodilia classes, peaked in testudines. However, across this period in time; CSF1R/CSF1 and CSF3R/CSF3 had become established as blood myeloid cell-specific factors and were unlikely to be driving the changes in WBC distribution (Figure 4bi). Notably, as the neutrophil populations declined, both the basophil and eosinophil populations increased, with basophils peaking in the Crocodilia lineage before a dramatic decline in testudines. Suggesting, environmental pressure on the immune response of certain orders favoured either the basophil or the eosinophil at the expense of neutrophils and the other granulocyte (Figure 4bi). The decline was not only restricted to neutrophil population, as the monocyte population had a similar pattern. Monocytes were at their highest levels in squamata before a rapid decline in crocodilia and becoming almost completely absent from Testudines. At the same time, the azurophil emerged effectively taking over the role of the monocytes in crocodilia and testudines (Figure 4bi).

Interestingly, by the advent of testudines, neutrophil/heterophils are no longer the dominant granulocyte, although granulocytes overall still represent the largest proportion of myeloid WBC (Figure 4bii). However, as the orders of aves, monotremata and marsupalia emerge there is a noticeable change. A limitation of visualising data in this form is the implication of the distinct sequential evolution of different animal orders. However; it’s likely there would have been a degree of overlap between the emergence of distinct orders. Therefore, it is noticeable that there was a rapid increase in the both avian heterophil population and monotremata neutrophil population compared to testudines (Figure 4bii). The increase in neutrophils was mirrored by an equally rapid decrease in eosinophils in both Aves and the mammalian orders (Figure 4bii). Neutrophils and eosinophils also share early common progenitors as part of the development pathway in the bone marrow, therefore neutrophil production appeared to be dominating the cell machinery. Interestingly, the switch to the predominance of the neutrophil over the eosinophil, coincides with the emergence of endothermy, strongly suggesting that external factors are behind these changes in myeloid WBC distribution as there are no noticeable changes in the distribution of the CSF1R/CSF1 and CSF3R/CSF3 ligand/receptor pairings. However, the impact is filtering down to the gene/protein level as there are changes in the shared sequence similarity of CSF3, as it increases more rapidly between Testudines and Monotremata, than in the period between Squamata and Testudines. Interestingly, these changes were not reflected in the CSF3R trajectory (Figure 4bii) Presumably, the arrival of endothermy resulted in the appearance of novel pathogens for which a neutrophil-mediated response was more appropriate than an eosinophil-mediated one, leading to reductions in the eosinophil population and virtual disappearance of basophils (Figure 4bii). The monocyte population had re-emerged as a single population by the appearance of aves and the distribution remained consistent in both the avian and mammalian orders (Figure 4bii). Intriguingly, the greatest period of change for myeloid WBC distribution is between testudines and monotremata. However, the equivalent period for protein homology happens much later between the mammalian orders of marsupalia and placentalia (Figure 4bii) and is common across all CSFR/CSF pairings. Therefore, as WBC function and distribution had previously been established in early lineages, this suggested that another external factor was responsible for the rapid change, possibly the emergence of internalized pregnancy (Figure 4bii).

## Discussion

The mammalian neutrophil is a highly specialised cell that acts as a first responder to insults against a host immune system as well as acting in an equally important sentinel role [2, 3]. They are functionally conserved across phylum Chordata and constitute the largest population of myeloid cells in birds and mammals. In humans, for example, up to one billion neutrophils per kilogram of body weight are produced in the bone marrow each day [4]. The immune response has evolved in such a way as to be able to respond efficiently to a variety of threats, and it is interesting that a resource such as the bone marrow should be expended on the production and maintenance of the relatively short lived neutrophil at the expense of the eosinophil and basophil. The timescale modelling demonstrates that prior to the advent of tetrapoda lineages, the neutrophil was in a pool of different WBC populations at the disposal of the jawed/jawless fish. However, the appearance of CSF3, altered the distribution and with each emerging animal order, a different granulocyte was favoured, presumably, for cell-specific adaptations in response to environmental challenges. Thus, suggesting that CSFR1R/CSF1 and CSF3R/CSF3 signalling conferred adaptions on the neutrophil that proved evolutionarily advantageous.

The bone marrow is an essential site for granulopoiesis and myelopoiesis. As multipotent haematopoietic stem cells (HSCs) progress through different vascular niches within the bone marrow they sequentially lose their potential to form other lineages in response to environmental cues from bone marrow dwelling macrophages and stromal endothelial cells [3, 4, 29]. The emergence of CSF1 in tetrapod lineages, that evolved after Jawed/Jawless fish, lead to the appearance of bone and bone marrow. A consequence of this was the gradual re-organisation of haematopoiesis away from existing haematopoietic tissues, such as the eosinophil rich Leydig organ of sharks to tissue-specific compartments in the bone marrow. CSF1, in conjunction with another factor, receptor activator of nuclear factor-κB ligand (RANKL), co-ordinate the reabsorption of old bone through haematopoietically derived osteoclasts to allow the generation of new bone. CSF1 and RANKL (which emerged at a similar point in evolution) orchestrate bone-remodelling through their respective receptors, CSF1R and RANK [30]. This suggests that by the time a bone structure had evolved in amphibia, the bone marrow had become the principle site of haematopoiesis, although they are some exceptions within various anurans [31].

In contrast to peripheral blood, CSF3 is expressed on a number of cells in the bone marrow including; neutrophils, monocytes, B cells, myeloid progenitors and HSCs [29, 32–34]. Although CSF3 can likely act directly on HSCs through their receptor, it is believed to indirectly mobilise HSCs through a monocytic intermediary that secretes CSF3, which leads to suppression of the CXCR4:CXCL12 axis, alteration of the bone marrow niche, and the subsequent release of HSCs [29, 32–34]. In a similar process, CSF3 can suppress B cell lymphopoiesis by again targeting CXCL12 and suppressing other B cell tropic factors or stromal cells that favour the lymphoid niche [35]. Thus, the emergence of CSF3R/CSF3 conferred the adaption, or advantage, of control of the biological machinery i.e. CSF3 provides a mechanism through which haematopoiesis can be shaped and deployed in favour of maximal neutrophil generation. This broadly aligns with the neutrophil starting to dominate peripheral populations in the transition from Amphibia to the lizards of early Squamata following the appearance of bone.

Mammalian neutrophil production occurs in the haematopoietic cords present within the venous sinuses and the daily output is approximately 1.7×10^9^/kg [36]. As our data demonstrate high levels of neutrophil output are common across warm-blooded chordates and neutrophil/heterophils are the predominant granulocyte in birds and mammals, in what could be considered an example of convergent evolution. One of the fundamental requirements of the host innate immune response is to be able to respond rapidly to perceived threats that can be present at any site in the body, which requires the cellular arm to be constantly present and available. Theoretically this can be achieved in the steady state by having either low volumes of long-lived cells or high volumes of short-lived cells circulating in the periphery. As a consequence, there are potential biological trade-offs when considering each setting, firstly; the longer a cell survives in a periphery, the more effort is required by the host to maintain its survival in terms of providing appropriate cues and growth factors. Secondly, while a short-lived cell does not need as much host input for survival, a high turnover is required in order to ensure it doesn’t compete for growth factors with other cell types or cause damage by being retained beyond its usefulness. The latter setting fits with the observed neutrophil life cycle and could be considered an evolutionary adaption. The CSF3R/CSF3 signalling pathway is essential for generating high neutrophil numbers without adverse effects to the host.

The neutrophil has a short circulatory half-life in the steady state of approximately one day [3, 4], although this does remain controversial as some estimates of the circulatory lifespan are as much as five days [37]. By contrast, a mature eosinophil has a short circulatory lifespan of approximately 18 hours before relocating to the tissue, where it will survive for a further 2-5 days [38]. Similarly, intermediate and non-classical monocytes can survive in the periphery for four and seven days respectively [39]. In human studies, CSF3 has been shown to “effectively” shorten the lifespan of neutrophil myeloblasts and promyelocytes by decreasing their time spent in cell cycle and accelerating their progress to maturation [40]. Although, it is unclear what the exact effect is on the lifespan of post-mitotic neutrophils, studies have shown that the addition of exogenous of CSF3 can delay apoptosis in mature neutrophils [41, 42]. This suggests that under certain conditions; such as infection, CSF3 can extend the lifespan of a mature neutrophil. Interestingly, classical monocytes also express CSF3R and have a short circulating lifespan of approximately 24 hours [39] These studies suggest that under steady state conditions and below a certain threshold, CSF3 has an as-yet-undocumented role either directly or indirectly in maintaining a short neutrophil lifespan and thus allowing the efficient turnover required to sustain high levels of granulopoiesis.

CSF3 orchestrates the life cycle of neutrophils in the bone marrow microenvironment by marshalling neutrophil progenitors through different development stages through to maturity. Accordingly, in response to this, CSF3R is expressed though every life stage, although at differing levels across the neutrophil populations, with the highest levels observed on mature cells where it is expressed at between two and three-fold more than on progenitors [43]. CSF3R/CSF3 signalling has a dual role in controlling the distribution of neutrophils by both retaining a population of mature neutrophils in the bone marrow as a reservoir and facilitating the egress of other neutrophils into the periphery. Thus, a major evolutionary adaptation that CSF3R/CSF3 has conferred to the neutrophil is the ability to move, both on a population-wide and individual cell level.

There is abundant production of neutrophils daily in the bone marrow; however, there are far fewer neutrophils in circulation in the blood than are produced during granulopoiesis as the total neutrophil population is effectively stored in the bone marrow or marginated in intravascular pools within the spleen and liver [44]. Neutrophils can transit between sites in response to CSF3 signalling and it is estimated that 49% of cells are present in the circulating pool and the remaining 51% are marginated in discrete vascular pools [36, 45]. In the event of an infection, neutrophils can be mobilised from the marginated pools and bone marrow in response to GCSF-induced production of mobilising signals such as CXCL1 [46]. The co-ordinated egress of neutrophils from the bone marrow is achieved by CSF3 interacting with the CXCR4/CXCL12 and CXCR2/CXCL2 signalling pathways. CSF3 disrupts the CXCR4-CXCL12 retention axis by reducing CXCL12 release from endothelial stromal cells and reducing CXCR4 expression on neutrophils, thus allowing the movement of mature neutrophils through the venous sinuses to the periphery. CSF3-induced expression of CXCR2 on neutrophils then causes their migration to the vasculature along a CXCL2 chemotactic gradient [1, 23, 24]. The limited study data presented here demonstrated that the antagonism between CXCR4 and CXCR2 is present *in vitro*, as in the absence of environmental cues from endothelial cells, exogenous CSF3 induced CXCR2 expression and reduced CXCR4. Interestingly, there was not a uniform impact as a minority of cells upregulated CXCR4 and downregulated CXCR2 as well CSF3R. Furthermore, CSF1R expression was subsequently induced on this minority population in a similar fashion observed to *ex vivo* cells. While this study is too small to draw firm conclusions, it potentially suggests there is an antagonistic interaction between CSF3R/CSF3 and CSF1R/CSF1 and work is ongoing to further define the role this antagonism plays in neutrophil biology.

As neutrophils enters the bloodstream, they need to be able to migrate easily around the circulatory system under high flow conditions. This requires the neutrophil’s physical form to have high deformability and flexibility as it encounters the different diameters of the vasculature. The neutrophil nucleus is functionally adapted to this role because of its multi-nucleated structure and the distinct protein composition of the nuclear envelope, features that are widely conserved across mammalian species [47]. CSF3 in co-ordination with C/EBPε and the ETS factors; Pu.1 and GA-binding protein (GABP) are responsible for the transcriptional control of the essential neutrophil nuclear structural proteins Lamin A, Lamin C and Lamin B receptor (LBR) [48, 49]. In comparison to other cell nuclear protein compositions, neutrophils have a low proportion of Lamin A and Lamin C, which is believed to make the nucleus more flexible for easier transit [47–49]. In contrast, the levels of LBR are increased, which is required for nuclear lobulation and subsequent neutrophil maturation [47, 48]. Therefore, CSF3 signalling and C/EBPε are intrinsic to the evolution of neutrophil cell motility. Intriguingly, the avian heterophil is not multi-lobulated, which aligns with our findings that C/EBPε had been lost from the lineage. This suggests that a different C/EBP family member is involved in the later stages of avian heterophil development.

The evolution of CSF3R/CSF3 has been essential to the development of the neutrophil/heterophil in chordates and its own existence as the principle neutrophil growth factor is evolutionarily advantageous. The Jawed/jawless lineages are unique in that they have populations of both neutrophils and heterophils, whereas later linages favour either neutrophils or heterophils. Our analysis shows that CSF3R/CSF3 emerged before the advent of Jawed/jawless fish as both are present in the Coelacanth and absent from the other lineages. Intriguingly, an IL6-like gene was present in the same syntenic location of the whale shark, suggesting the possibility that heterophils and neutrophils were independently controlled by an IL6-like protein and CSF3 in the early lineages. IL6 is an important pleiotropic pro-inflammatory cytokine that plays key roles in infection, inflammation and haematopoiesis [50]. Although CSF3R/CSFR and IL6/IL6R, which are functional paralogues, diverged from each other many millions of years ago, there is still functional redundancy at the cell level in modern-day mammals, as neutrophils are present - though at vastly reduced numbers - in CSF3^−/−^ and CSF3R^−/−^ mice [10, 11]. IL-6 can act on neutrophil progenitors and immature neutrophils thus supporting neutrophil development in the bone marrow of CSF3 deficient mice [12]. However mature neutrophils are refractory to IL-6 signalling as the expression of the IL-6R subunit gp130 is lost during maturation [51]. From amphibia onwards, the protein analysis here suggests that CSF3R and CSF3 co-evolved closely together and in contrast to CSF1R/CSF1/IL34, CSF3 is likely the only ligand of CSF3R.

Our findings have demonstrated how essential CSF3R and CSF3 (and to a lesser extent CSF1R and CSF1) are to the predominance of mammalian neutrophils in blood. Through the course of evolution, CSF3R/CSF3 signalling has accrued many properties that are responsible for the survival of the neutrophil and its ability to function, such as cell motility and mobility. Interestingly, although CSF3 is present in jawed/jawless fish lineages, it’s not until the emergence of tetrapoda that CSF3R and CSF3 begin to dominate haematopoiesis. Our current studies have been limited to chordates in which the data is publicly available. However, we envisage exploring the phagocytic granulocyte of two model species. The amphioxus is considered the basal chordate and a macrophage-like population has been identified, although a neutrophil/heterophil has yet to be described within the rudimentary circulatory system [52]. The lungfish, a primitive airbreathing fish, is unique among jawed/jawless lineages as it lives in freshwater and can survive on the land for up to one year. Accordingly, the lungfish has many adaptations and is considered to be a one of the closest living relatives to tetrapods making it a good model species for further study [53]. Given how essential motility and mobility are to neutrophil function and development, it would be useful to discern when it emerged in evolution by identifying if equivalent cells and functional gene/protein orthologues are present in either species. These comparative studies would answer fundamental questions about the origin of the neutrophilic phagocyte.

## Materials and Methods

### Species selection

Ninety-seven animals from all the major classes were selected, thirty-eight of which, were used for the calculation of haematological parameters and the remaining fifty-nine for the bioinformatics-based studies. Analyses was performed on non-mammalian lineages including; jawed/jawless fish, amphibia, non-avian reptiles – both non-crocodilian and crocodilian-, aves, and the mammalian lineages; monotremata, marsupialia and placentalia. Urochordates (tunicata), and cephalochordate (Amphoxi) were excluded from analysis as there was insufficient coverage or insufficient annotation of sequence data in the Pubmed Gene and Ensembl databases. Similarly, the lungfish (Dipnoi) was also excluded from this analysis because of insufficient coverage of sequence data. Finally, teleost lineages were excluded from the analysis owing to having gone through three rounds of genome wide duplication, in contrast to all other chordates who have only undergone two rounds [54]. A species tree was generated using the NCBI Common Taxonmy browser common tree tool [55, 56] and visualised using the Interactive Tree of Life (iTOL) web browser tool [57].

### Comparative analysis of haematological parameters and CSF/CSFR gene presence and synteny in chordates

Complete blood count data for thirty-eight species representing each animal order or class were collated from existing literature to perform a meta-analysis of myeloid WBC in phylum Chordata. Only counts that were calculated to the standardised concentration (10^9^/L) or equivalent were used and all lymphoid data was excluded. The proportional composition for each myeloid subset; neutrophil, eosinophil, basophil and monocytes as part of the overall myeloid population was also calculated.

Gene sequence for fifty-nine species - representing different animal orders and classes - were retrieved from the NCBI Gene databases [58] in most instances or from annotated entries in Ensembl (v95) [59] for the following genes; CSF1R, IL34, CSF1, CSF3R, CSF3, C/EBPα, C/EBPβ C/EBPδ and C/EBPε. A heatmap visualising gene presence, absence or presence of orthologues was generated using maplotlib/seaborn library for Python 3.7.7.

Syntenic maps were generated manually for selected genes in the Jawed fish lineage. To generate the maps, syntenic blocks were identified as a section of the human chromosome containing the gene of interest flanked by *x* number of genes. The blocks were then manually compared to similar regions in the chromosomes in Jawed fish to identify orthologous genes. Between three and five genes were chosen per syntenic block to function as anchor points of reference. Anchor points were selected based on their situational proximity, either upstream or downstream, to gene of interest and were present in all species examined. Multiple genes were identified as anchor points to mitigate for the random loss of genes during the process of evolutionary gene rearrangement.

### Generation of percentage shared sequence similarity plots and timescales

As with the gene sequences, protein sequences for the identified species were retrieved from the NCBI Protein databases. Presumed orthologous sequences were screened using the NCBI Basic Local Alignment Search tool (BLAST) to generate a percentage score of sequence similarity. The parameters to determine shared sequence similarity were as follows; a sequence was deemed to be homologous where coverage of the protein sequence was equal to or greater than 40% of the total protein sequence examined and the e value was between 0×10^−20^ - 0. Percentage scores were averaged per order. Shared sequence similarity was also plotted against animal order or class. The same data was also used to generate two sets of timescales for each order/ member for a given receptor/ligand family, where either the calculated mean shared sequence similarity value or haematological parameters were plotted on a timescale based on the emergence of the earliest known ancestor of that order *versus* time (million years ago [mya]).

### Isolation of peripheral blood myeloid cells and cell culture

Peripheral blood mononuclear cells (PBMCs) and Polymorphonuclear cells (PMNs) were isolated from the blood of healthy donors using a triple density Percoll™ gradient. Both cell fractions were washed and incubated in isotonic buffer solution for downstream applications. Monocytes and neutrophils were harvested from the PBMC and PMN fractions using immunomagnetic negative and positive selection respectively and the protocols were performed according to the manufacturer’s recommendation (Stemcell Technologies, Cambridge, UK).

#### Mobility assays

Isolated monocytes were incubated in either CSF3 (20ng/ml) or CSF1 (20ng/ml) for 72 hours prior to the set-up of the assay. The CSF-treated cells were recovered and washed in isotonic solution before being loaded into an Ibidi 2D chemotaxis chamber to assess chemotaxis in response to a serum-based gradient. Migration velocity was tracked by microscopy for one hour following seeding of cells. A parallel control stream using freshly isolated neutrophils was also set up.

#### CXCR4 induction

Isolated neutrophils were cultured *in vitro* for 20 hours in 10% Foetal Bovine Serum (FBS) supplemented RPMI 1640 culture medium to induce CXCR4 expression. A separate control culture of neutrophils with the addition of exogenous CSF3 (20ng/ml) was also established. At the conclusion of the incubation period, both cell cultures were recovered, washed in isotonic solution and used for downstream analytical flow cytometry.

### Flow cytometry

All antibodies used were purchased from Biolegend, London, UK. Freshly isolated PBMCs or PMNs or cultured neutrophils were incubated with various antibodies as per manufacturer’s instructions for cell surface staining. For the identification of CSF3R and CSF1R expression, cells were stained with fluorophore conjugated monoclonal anti-human antibodies against CD3 (OKT3), CD19(HIB19), CD56(39D5), NKp46 (9E2) [lin^−^], CD14 (63D3), CD16 (3G8), CD66b (G10F5), CSF3R (LMM741), CSF1R (9-4D2-1E4) and HLA-DR (L2434). After exclusion of cell debris and doublets based on side scatter characteristics (SSC), monocytes were identified as SSC^int^ lin^−^ CD14^+^CD16^+/-^HLA-DR^+^. The granulocyte population was identified as SSC^hi^ Lin^−^CD66b^+^CD16^hi^HLA-DR^+^ and the minority eosinophil population was excluded from neutrophils by differential CD16 expression. To further characterise CXCR4 induced neutrophils, the following monoclonal antibodies were also incorporated CXCR4 (12G5) and CXCR2 (5E8/CXCR2).

