## Supplementary Figure 1 for "Analysis of receptor-ligand pairings and distribution of myeloid subpopulations across the animal kingdom reveals neutrophil evolution was facilitated by colony-stimulating factors" (from another version of this preprint: version 1, 2020-06-20)

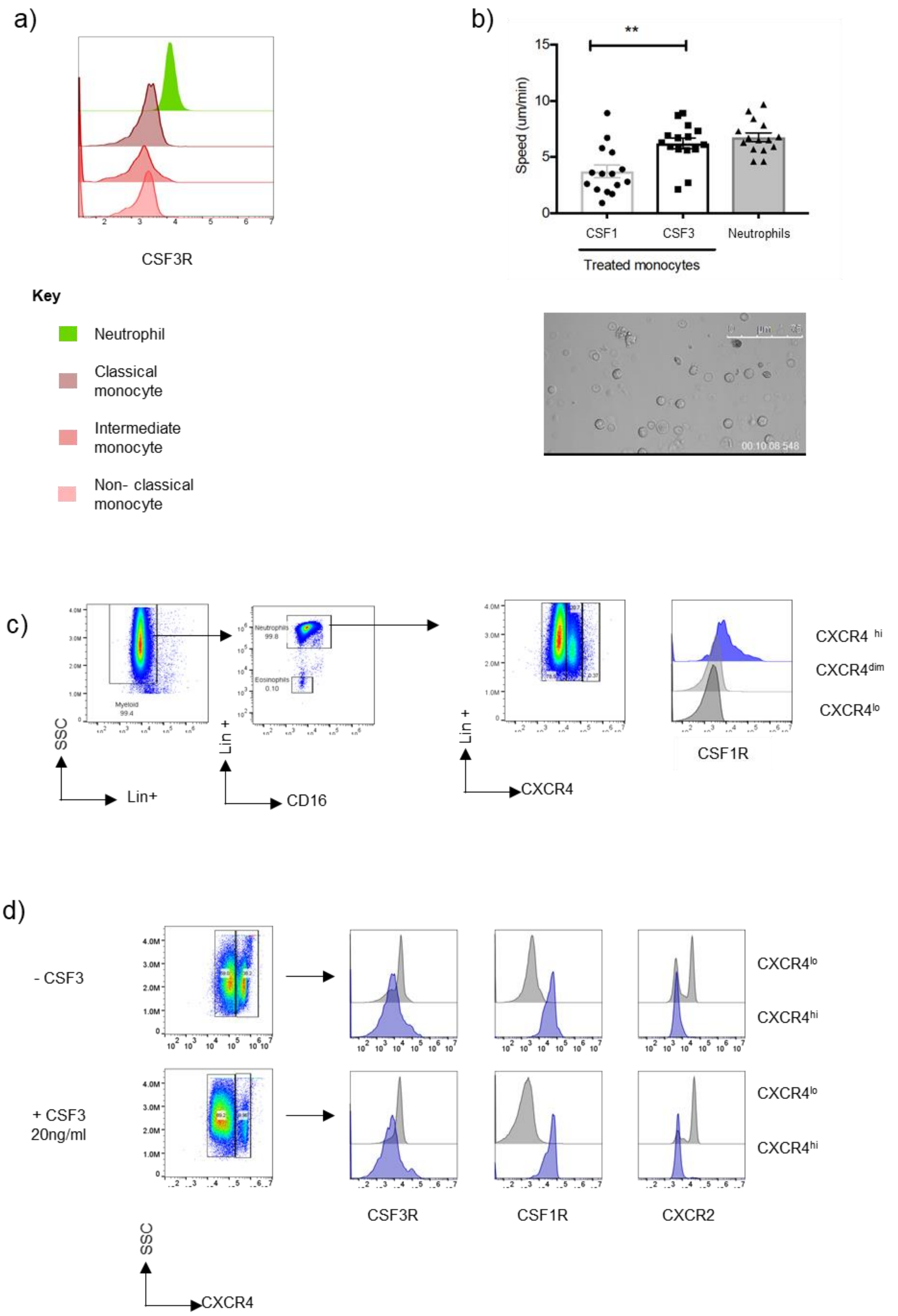
a)
CSF3R
Key
Neutrophil
Classical monocyte
Intermediate monocyte
Non- classical monocyte
b)
**
15
10
5
0
Speed (um/min)
CSF1
CSF3
Neutrophils
Treated monocytes
00:10:08.548
c)
Myeloid
99.4
Neutrophils
99.8
Eosinophils
0.10
SSC
Lin+
Lin +
CD16
Lin +
CXCR4
CSF1R
CXCR4 hi
CXCR4dim
CXCR4lo
d)
- CSF3
+ CSF3
20ng/ml
CXCR4lo
CXCR4hi
CXCR4lo
CXCR4hi
CSF3R
CSF1R
CXCR2
SSC
CXCR4

**Supplementary figure 1: Cross expression and potential functionality of CSF1R and CSF3R in human monocytes and neutrophils *in vitro***

A. Representative plot of differential CSF3R expression in monocyte subsets and neutrophils. B. Velocity plot of 72 hour CSF1-treated monocytes, CSF3-treated monocytes and freshly isolated neutrophils. C. Representative plot of CSF3R expression in CXCR4 subsets of freshly isolated neutrophils. D. Representative plots of CSF3R expression in induced $CXCR4^{hi}$ and $CXCR4^{lo}$ neutrophil populations following 24-hour culture in the presence of absence of exogenous CSF3 (20ng/ml)
